# Knock-in rat lines with Cre recombinase at the dopamine D1 and adenosine 2a receptor loci

**DOI:** 10.1101/455337

**Authors:** Jeffrey R. Pettibone, Jai Y. Yu, Rifka C. Derman, Thomas W. Faust, Elizabeth D. Hughes, Wanda E. Filipiak, Thomas L. Saunders, Carrie R. Ferrario, Joshua D. Berke

## Abstract

Genetically-modified mice have become standard tools in neuroscience research. Our understanding of the basal ganglia in particular has been greatly assisted by BAC mutants with selective transgene expression in striatal neurons forming the direct or indirect pathways. However, for more sophisticated behavioral tasks and larger intracranial implants, rat models are preferred. Furthermore, BAC lines can show variable expression patterns depending upon genomic insertion site. We therefore used CRISPR/Cas9 to generate two novel knock-in rat lines specifically encoding Cre recombinase immediately after the dopamine D1 receptor (*Drd1a*) or adenosine 2a receptor (*Adora2a*) loci. Here we validate these lines using *in situ* hybridization and viral vector mediated transfection to demonstrate selective, functional Cre expression in the striatal direct and indirect pathways respectively. We used whole-genome sequencing to confirm the lack of off-target effects, and established that both rat lines have normal locomotor activity and learning in simple instrumental and Pavlovian tasks. We expect these new D1-Cre and A2a-Cre rat lines will be widely used to study both normal brain functions and neurological and psychiatric pathophysiology.

## INTRODUCTION

Dopamine and adenosine are important chemical messengers in the brain, vasculature, and elsewhere in the body. Within the brain, one key site of action is the striatum, a critical component of basal ganglia circuitry involved in movement, motivation, and reinforcement-driven learning (Berke, 2018; Gerfen and Surmeier, 2011). Most (90-95%) striatal neurons are GABAergic medium spiny neurons (MSNs) with two distinct subclasses (Gerfen and Surmeier, 2011). “Direct pathway” neurons (dMSNs) express dopamine-D1 receptors and project primarily to the substantia nigra pars reticulata / globus pallidus pars interna (SNr/GPi), whereas “indirect pathway” neurons (iMSNs) express both dopamine-D2 receptors and adenosine-A2a receptors, and project primarily to the globus pallidus pars externa (GPe). Although our understanding of their distinct functions is incomplete, dMSNs and iMSNs have complementary roles promoting and discouraging motivated behaviors respectively (Collins and Frank, 2014).

The investigation of dMSNs and iMSNs has been transformed by transgenic mice. Random genomic insertion of BACs (bacterial artificial chromosomes) encoding dopamine receptor promoters driving fluorescent protein expression confirmed the near-total segregation of striatal D1 and D2 receptors (Matamales et al., 2009; Shuen et al., 2008) and enabled identification of dMSNs/iMSNs in brain slices (Day et al., 2006). BAC lines in which dopamine receptor promoters drive Cre recombinase expression (D1-Cre, D2-Cre, etc) have allowed *in vivo* identification and manipulation of neuronal subpopulations in striatum (Barbera et al., 2016; Cui et al., 2013; Kravitz et al., 2010, 2012) and cortex (Kim et al., 2017). iMSNs targeting is further improved using an A2a promoter, rather than D2, because A2a receptors are selectively expressed on iMSNs while D2 receptors are also expressed on other striatal cells and synapses (Alcantara et al., 2003).

However, for many experiments rats are more suitable than mice. Their larger size means they can bear complex intracranial implants without loss of mobility. Furthermore, rats can learn more sophisticated behavioral tasks, including those investigating reinforcement learning (Hamid *et al*, 2016) and behavioral inhibition (Schmidt et al., 2013). The advent of CRISPR/Cas9 methods has facilitated the generation of knock-in rat lines (Jung et al., 2016; Mali et al., 2013), and knock-ins are more likely to have faithful expression patterns compared to BACs for which (for example) different D1-Cre lines show markedly different expression (Heintz, 2004).

Here we describe the generation of transgenic D1-Cre and A2a-Cre rat lines using CRISPR/Cas9, and then demonstrate the specificity of *iCre* mRNA expression in the intended striatal targets. Next, we confirm Cre-dependent expression to demonstrate that Cre is functional and appropriately confined to the direct or indirect pathways. Finally, we demonstrate normal locomotor activity, learning and motivation in simple behavioral tasks.

## MATERIALS AND METHODS

All animal procedures were approved by the University of Michigan and UCSF Institutional Animal Care & Use Committees.

### Genetic Engineering

CRISPR/Cas9 (Cong et al, 2013; Mali et al, 2013) was used to generate genetically-modified rat strains. Two single guide RNA (sgRNA) targets and protospacer adjacent motifs (PAM) were identified downstream of the rat *Adora2a* termination codon (Hsu et al., 2013). sgRNA targets were cloned into plasmid pX330 (Addgene.org #42230, a gift of Feng Zhang) as described (Ran et al., 2013). Guide targets were C30G1: CTAAGGGAAGAGAAACCCAA PAM: TGG and C30G2: GGCTGGACCAATCTCACTAA PAM: GGG. Purified pX330 plasmids were coelectroporated into rat embryonic fibroblasts with a PGKpuro plasmid (McBurney et al., 1994). Genomic DNA was prepared after transient selection with puromycin (2 μg/ml). A 324bp DNA fragment spanning the expected Cas9 cut sites was PCR-amplified with forward primer GGGATGTGGAGCTTCCTACC and reverse primer GCAGCCCTGACCTAACACAG. DNA sequencing of the amplicons showed that C30G1-treated – but not C30G2-treated - cells contained overlapping chromatogram peaks, indicative of multiple templates that differ because of non-homologous end-joining repair of CRISR/Cas9-induced chromosome breaks resulting in the presence of small deletions/insertions (indels). sgRNA C30G1 was chosen for rat zygote microinjection. A DNA donor was synthesized (BioBasic.com, cloned in pUC57) to introduce the following elements between codon 410 and the termination codon of *Adora2a*: a glycine-serine-serine linker with porcine teschovirus-1 self-cleaving peptide 2A (P2A; (Kim et al., 2011)) followed by iCre recombinase (Shimshek et al., 2002) with hemagglutinin tag YPYDVPDYA (Kolodziej and Young, 1991) and a termination codon with the bovine growth hormone polyadenylation sequence (Goodwin and Rottman, 1992). To mediate homologous recombination a 5’ arm of homology (1804bp of genomic DNA 5’ to codon 410) and a 3’ arm of homology (1424bp of genomic DNA downstream of the termination codon) were used. The 20bp sequence of C30G1 was omitted from the 3’ arm of homology to prevent CRISPR/Cas9 cleavage of the chromosome after insertion of the DNA donor.

A similar approach was used for *Drd1a*. Two sgRNA were identified downstream of the *Drd1a* termination codon - C31G1: TTCCTTAACAGCAAGCCCAA PAM: GGG and C31G2: CTGAGGCCACGAGTTCCCTT PAM: GGG. A 293bp DNA fragment spanning expected Cas9 cut sites was PCR-amplified with forward primer TGGAATAGCTAAGCCACTGGA and reverse primer CTCCCAAACTGATTTCAGAGC. Both sgRNAs were found to be active after transfection in rat fibroblasts by T7 endonuclease 1 (T7E1) assays (Sakurai et al., 2014). Briefly, DNA amplicons were melted and re-annealed, then subjected to T7EI digestion. The presence of indels produced by non-homologous endjoining repair of Cas9-induced double strand breaks resulted in the presence of lower molecular weight DNA fragments for both sgRNA targets, and C31G1 was chosen for zygote microinjection. A DNA donor was synthesized (BioBasic.com, cloned in pUC57) to introduce the following elements between *Drd1a* codon 446 and the termination codon: a glycine-serine-serine linker with P2A followed by *iCre* recombinase with V5 peptide tag GKPIPNPLLGLDST (Yang et al., 2013) and a termination codon with the bovine growth hormone polyadenylation sequence. To mediate homologous recombination a 5’ arm of homology (1805bp of genomic DNA 5’ of codon 446) and a 3’ arm (1801bp of genomic DNA downstream of the termination codon) were used. The 20bp sequence of C31G1 was omitted from the 3’ arm of homology to prevent cleavage of the chromosome after insertion.

**Rat Zygote Microinjection** was carried out as described (Filipiak and Saunders, 2006). sgRNA molecules from a PCR-amplifìed template were obtained by *in vitro* transcription (MAXIscript T7 Transcription Kit followed by MEGAclear Transcription Clean-Up Kit - Thermo Fisher Scientific). The template was produced from overlapping long primers (IDTDNA.com) that included one gene-specific sgRNA target and T7 promoter sequence that were annealed to a long primer containing the sgRNA scaffold sequence (Lin et al., 2014). Cas9 mRNA was obtained from Sigma-Aldrich. Circular DNA donor plasmids were purified with an endotoxin-free kit (Qiagen).

Knockin rats were produced by microinjection of a solution containing 5 ng/μl Cas9 mRNA, 2.5 ng/μl sgRNA, and 10 ng/μl of circular donor plasmid. Prior to rat zygote microinjection fertilized mouse eggs were microinjected with the nucleic acid mixtures to ensure that the plasmid DNA mixtures did not cause zygote death or block development to the blastocyst stage. Rat zygotes for microinjection were obtained by mating superovulated Long Evans female rats with Long Evans male rats from an inhouse breeding colony. 353 rat zygotes were microinjected with A2a-Cre reagents, 289 survived and were transferred to pseudopregnant SD female rats (Strain 400, Charles River), resulting in 60 rat pups. 401 rat zygotes were microinjected with D1-Cre reagents, 347 survived and were transferred, resulting in 95 pups. Genomic DNA was purified from tail tip biopsies (Qiagen DNeasy kit) to screen potential founders for correct insertion of *iCre*.

### G0 Founder Screening Primers

*iCre* internal primers for both lines:

FWD AATGTGAACATTGTGATGAACTACA, REV

CAGAATAGAATGACACCTACTCAGACA

Insertion-spanning primers for A2a-Cre:

5’ Junction FWD: AGGCAACTTTCTAGTTGACAAATCAAG,

REV: CAGCAGGCTGAAGTTAGTAGCTC

3’ Junction FWD: CATTGTCTGAGTAGGTGTCATTCTATTCT,

REV: GAATCACAGCCCAAGAGATACTACACT

Insertion-spanning primers for D1-Cre:

5’ Junction FWD: AAAAGTGACTAGAATTGACCTGGAAGAG, REV:

AGCAGGTTGGAGACTTTCCTCTTCTTCTT

3’ Junction FWD: CATTGTCTGAGTAGGTGTCATTCTATTCT,

REV: GGAAAAGGAAAGAGAAGCAGAATAAT.

### Colony management and genotyping

Lines were maintained by backcrossing with wild-type Long Evans rats (Charles River or Harlan). Offspring were genotyped using real-time PCR (Transnetyx), using the following insertion-spanning primers:

A2a-Cre: FWD: CGTCTCCAGCCTGCTTCAG, REV: TCCTCATGGTCTTCAGAGTTTGC

Reporter 1: CCGGAAGCGGAGCTAC

D1-Cre: FWD: GTGAGGCTGCTCGAGGAT, REV: CTGGCAACTAGAAGGCACAGT

Reporter 1: CCTGGACAGCACCTGAC

**Genome sequencing was performed using** blood samples (1 mL per rat) at the UCSF Institute for Human Genetics. Libraries were prepared from fragmented DNA (Kapa Hyper Prep) and sequenced (Illumina NovaSeq 6000, S4 flow cell, paired-end mode, read lengths 150bp). Sequencing reads were aligned to the rat genome (RGSC Rnor_6.0) using the Burrows-Wheeler Aligner (BWA-MEM). We used GATK HaplotypeCaller, Samtools, Bedtools, Pysam and Matlab for variant calling, subsequent analysis and visualization.

To determine the location of the inserted *iCre* cassette, we selected reads that did not align as a pair to the rat genome, which includes reads where only one mate or no mate of the pair aligned to the genome. These unaligned reads will include matches to the inserted *iCre* cassette sequence, which is not part of the reference genome. We searched for paired-end reads where one mate is aligned to the *iCre* cassette, and then examined where in the genome the other mate is aligned.

To further verify the integrity of our lines we examined potential off-targets (D1-Cre: 197 sites and A2a-Cre: 557 sites) predicted by an *in silico* sgRNA off-target prediction algorithm (<u>CRISPOR.net</u> RSGC Rnor_6.0). CRISPR/Cas9-induced mutations in exons are of particular concern. Only two potential off-targets were predicted to be in exons in the D1-Cre line and none in the A2a-Cre line. After analyzing assembled genomic sequence data, we found the D1-Cre sequence contained a one base pair deletion at chr18:49935989 (Zfp608) and a single nucleotide variant (SNV) at ch1:258074844 (Cyp2c). Upon inspection these changes were present in both the D1-Cre line and the A2a-Cre line, consistent with natural variations in the Long Evans strain rather than off-target mutations from the D1-targeting sgRNA. Among the 195 predicted intronic off-targets for the D1-Cre strain located in introns, we observed 11 changes (8 contained SNVs and 3 contained indels). For the 557 predicted intronic off-target locations in the A2a-Cre strain, 48 locations showed changes (39 contained SNVs, 6 contained indels, and 3 contained both). Closer inspection of the indels revealed that 100% were present in both D1-Cre and A2a-Cre lines. This is again consistent with Long Evans strain variation rather than off-target changes.

### *In situ* hybridization

Frozen brains (n=6, one male and two females from each line) were stored at −80C (~overnight-2 weeks), then sectioned on a cryostat at 20 μm and mounted on glass slides. Sections were fixed in 4% PFA at 4C for 15 minutes and dehydrated through 50%, 75%, 100% and fresh 100% EtOH at RT for 5 minutes each. Slides were dried completely for 5 minutes. A hydrophobic barrier (Advanced Cell Diagnostics) was drawn around each section. Slides were rinsed twice in 1xPBS (~1-3 minutes) and incubated with Protease IV reagent (Advanced Cell Diagnostics) for 30 minutes at RT. Fluorescent probes (RNAScope, Advanced Cell Diagnostics, *iCre* Cat. 312281, *Drd1a* Cat. 317031-C2 and *Adora2a* Cat. 450471-C3) were added (2 hr, 40C) followed by manufacturer-specified washing and amplification. DAPI was added to the slides before coverslipping (Prolong Gold, Thermo Fisher Scientific).

We used MIPAR software (www.mipar.us) to segment cell boundaries and fluorescent puncta using separate processing pipelines. To define nuclear boundaries, the DAPI channel of each image was first histogram-equalized to compensate for uneven illumination (512×512 pixel tiles) and convolved with a pixel-wise adaptive low-pass Wiener filter (5×5 pixel neighborhood size) to reduce noise. The image was then contrast-adjusted (saturating the top and bottom 1% of intensities). Bright objects were segmented using an adaptive threshold (pixel intensity >110% of mean in the surrounding 30-pixel window). Image erosion followed by dilation further reduced noise (5-pixel connectivity threshold, 10 iterations). The Watershed algorithm was applied to improve object separation. Objects >5000 pixels (i.e. clustered nuclei) were identified and reprocessed to improve separation. Since mRNA fluorescent puncta can be located in the endoplasmic reticulum, we dilated the boundaries of each segmented nucleus by 5 pixels to include these regions.

To segment fluorescent puncta, each of the 3 probe channels were first preprocessed using a Top-hat filter (9-15-pixel radius), Wiener filter (15×15 pixel neighborhood size) followed by contrast adjustment (saturating top and bottom 1% of intensities). Bright regions were segmented using the extended-maxima transform (8-connected neighborhood, 5 H-maxima). A Watershed algorithm followed by erosion was used to improve object separation. Objects < 5 pixels were rejected as noise. The location of each punctum is defined as the centroid of the segmented object.

For each fluorescent probe image channel, we counted the number of segmented puncta lying within a nuclear boundary. To determine the puncta threshold for specific versus non-specific probe hybridization, we estimated the “baseline” number of puncta expected per nucleus by chance from non-specific hybridization. We first calculated the puncta count per pixel for all puncta lying outside of cell nuclei and then multiplied this value by the number of pixels for each DAPI-labeled nucleus. This background puncta count was assumed to follow a Poisson distribution, and we defined our threshold for categorizing a cell as “positive” for a given mRNA probe as the 95^th^ percentile of this distribution. Consistency was calculated as the percentage of *Drd1a*+ (or *Adora2a*+, in the case of A2a-Cre) nuclei that are also positive for *iCre*. Specificity was calculated as the percentage of *iCre*+ nuclei that are also *Drd1a*+ (or *Adora2a*+). Off-target consistency and specificity were calculated the same way, but substituting *Drd1a*+ (or *Adora2a*+) for each other in the above two equations.

### Virus injection

Rats (n=2 females, one from each line) were microinjected with 0.5 μL of AAV5-CAG-Flex-TdTomato virus (UNC Vector Core) in dorsal striatum at three locations along a dorsal-ventral trajectory (AP: +1.5, ML: +2.2, DV: −3.0, −4.0, −5.0 from brain surface), and killed 4 weeks after surgery.

### *In vivo* opto-tagging

Rats (n=2, one male from each line) were injected with 1.0 μL of hSyn-Flex-ChrimsonR-TdTomato virus (UNC vector core) bilaterally in ventral striatum (AP: +1.75, ML: +/−1.6, DV: −7.0 from brain surface) and implanted with two 64-channel drivable tetrode arrays, each with a fixed optical fiber extending centrally through the array to a depth of 6.5 mm. After 3 weeks of transfection, the tetrodes were lowered into the ventral striatum and recorded wideband (1-9000 Hz) at 30000 samples/s using an Intan digital headstage. Recording ended with a brief laser stimulation protocol (1 mW, 638 nm, 1-10 ms/ 1 Hz). The rat was awake, unrestrained, and resting quietly throughout the recording.

Units were isolated offline using automated spike sorting software (MountainSort, Chung *et al*, 2017) followed by manual inspection. For a unit to be considered a successfully-identified Cre+ neuron it had to meet several criteria: 1) evoked spiking within 10ms of laser onset, that reached the p< 0.001 significance level in the stimulus-associated latency test (Kvitsiani et al., 2013); 2) peak firing rate (Z-scored) of >10 during both 5 ms and 10 ms laser pulses; 3) a Pearson correlation coefficient >0.9 between their average light-evoked waveform and their average session-wide waveform.

### Imaging

Images were taken with a Nikon spinning disc confocal microscope with a 40X objective (Plan Apo Lambda NA 0.95). For viral tracing, images (2048×2048 pixels at 16 bit depth) were stitched in FIJI.

### Behavior

Rats were maintained on a reverse light-dark schedule (12/12), testing was conducted during the dark phase, and rats were at least 70 days old at the start of the studies. Males and females were used for instrumental and Pavlovian studies. Males were used to evaluate cocaine-induced locomotor activity because of well-established sex differences in response to cocaine (Becker and Koob, 2016) and an insufficient available number of females to examine them separately.

#### Instrumental and Pavlovian Procedures

Procedures were conducted in operant chambers as described (Derman and Ferrario, 2018). Rats (D1-Cre-, n=9 [3 males, 6 females]; D1-Cre+, n=16 [7m, 9f]; A2a-Cre-, n=8 [6m, 2f]; A2a-Cre+, n=16 [10m, 6f]). were food restricted to 85-90% of free-feeding body weight. For instrumental training, a food cup was flanked by two retractable levers. First, rats were given two sessions in which 20 food pellets (45 mg, Bioserv #F0021) were delivered into the food cup on a variable-interval schedule of 60s (VI60). Next, rats underwent instrumental training in which responses on the ‘active’ lever resulted in delivery of a single pellet (fixed ratio 1; FR1) and responses on the other ‘inactive’ lever had no consequences. Rats were trained to an acquisition criterion of 50 pellets within 40 min. The same rats then underwent Pavlovian conditioning using two auditory conditioned stimuli (CS; tone and white noise, 2 min; 4 presentations of each CS per session, 5 min ITI, 12 sessions, 1 hr/session). Fifteen seconds following CS+ onset, 4 pellets were delivered on a VI30 schedule. The CS-was presented an equal number of times, unpaired with pellets. Food cup entries were recorded in 10s bins and entries during the first 10s of CS presentations were used to evaluate conditioned anticipatory responding (i.e., prior to US delivery).

*Locomotor activity* was assessed in a subset of the rats trained above (Cre-, n=7; D1-Cre+, n=7; A2a-Cre+, n=7) using procedures similar to (Vollbrecht et al., 2016). Rats were allowed to feed freely for at least 5 days prior to locomotor testing. Testing was conducted in rectangular plastic chambers (25.4cm X 48.26cm X 20.32cm) outfitted with photocell arrays around the base perimeter. Beam breaks were measured using CrossBreak software (Synaptech; University of Michigan). Rats were habituated to the testing chambers (30 min) and given two injections of saline (1 mL/kg, i.p.) separated by 45 min. Next, the acute locomotor response to cocaine was assessed. After a 30 min habituation, rats were given a saline injection followed 45 min later by cocaine (15 mg/kg, i.p.) and remained in the chambers for an additional 60 min. Locomotor activity was recorded in 5 min bins throughout and reported as crossovers (beam break at one end of the cage followed by beam break at the opposite end of the cage).

## RESULTS

### Molecular Design

The D1-Cre and A2a-Cre rat lines were designed so that the native *Drd1a* or *Adora2a* promoter drives expression of both the native receptor and the codon-improved *Cre* recombinase (*iCre*) sequence in a single transcription event (Figure 1A). The use of iCre over Cre has been shown to enhance recombinase expression and limit epigenetic silencing in mammalian cells (Shimshek et al., 2002). For each line, a unique single strand guide RNA (sgRNA) was generated to induce double strand breaks at the terminus of the receptor coding sequence, and microinjected into Long Evans rat zygotes along with Cas9 and a circular plasmid containing the donor gene cassette. After correct recombination of the donor cassette, the 3’ end of target receptor sequence will be joined in frame with the “self-cleaving” peptide P2A (to separate the Cre protein after translation), followed by Cre with a nuclear localizing signal affixed at the amino terminus, and a peptide tag (HA for Adora2a, V5 for D1) to facilitate antibody-based detection.

**Figure 1.**
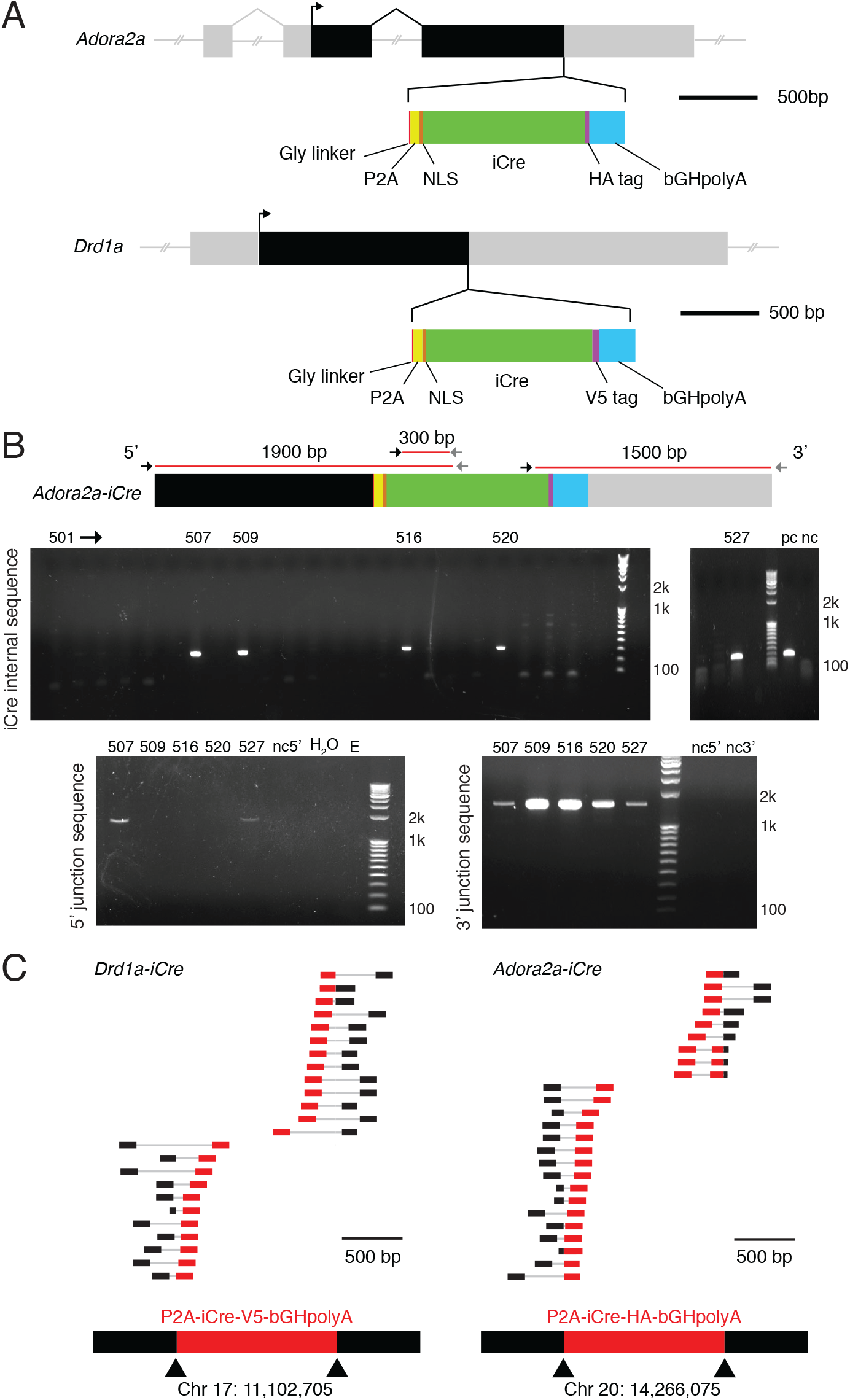
Details of insertion design and founder line screening. (A) Schematic of insertion cassettes into *Adora2a* (above) and *Drd1a* (below) genes. (Abbreviations: P2A, porcine teschovirus-1 self-cleaving peptide; NLS, nuclear localization sequence; HA, influenza hemagglutinin protein tag YPYDVPDYA; V5, peptide tag GKPIPNPLLGLDST; bGH, bovine growth hormone polyadenylation sequence). (B) PCR primer loci (above) and corresponding gels (below) demonstrating G0 screening of the A2a-Cre line. The top row of gels indicate that rats #507, 509, 516, 520 and 527 are transgenic for *iCre*. The bottom gels show that rats #520 and #527 have *iCre* inserted correctly at both the 3’ and 5’ junctions. See Methods for full primer sequences for screening both lines (Abbreviations: pc, single copy detection; nc, unrelated rat tail DNA; H2O, water control; E, empty). (C) Reads from whole genome sequencing aligned to a wild-type rat genome demonstrate that, for each transgenic line, the *iCre* cassette is inserted only once in the genome and at the target loci. Each row corresponds to one paired-end read, where one mate of the pair is aligned to the inserted cassette (red) and the other mate in the genome (black). Sequence reads with at least 100bp match in the inserted cassette are shown. All such pairs map to only one location in the genome.

### Founder screening, germline transmission and full genome sequencing

DNA samples from G0 potential founders were screened with primers to detect *iCre* in the genome (for primer sequences, see Methods). From this screen 21/96 potential D1-Cre, and 9/60 potential A2a-Cre founders were positive for *iCre*. Positive rats were then screened with additional primers across the junctions between native and introduced DNA stretches, to discriminate between correct and random genomic integration events (Figure 1B). This yielded 7/21 correct D1-Cre insertions and 7/9 correct A2a-Cre insertions. The *iCre* insert was then completely sequenced in these rats (14 total) to confirm complete integration.

These G0 founders were mated with wild-type Long Evans rats, and the G1 pups genotyped for *iCre* specific insertion as above to verify germline transmission. Colonies from one successful founder for each line were established and maintained by back-crossing to wild-type Long-Evans rats from commercial vendors (see Methods); all experimental results shown are from rats back-crossed for at least 3 generations

After 5 generations of back-crossing we took one female rat each from the D1-Cre and A2a-Cre lines and sequenced their entire genomes to confirm that *iCre* was present in the intended location and nowhere else (Figure 1C). Average sequencing depths for D1-Cre and A2a-Cre lines were 80x (1,503,983,138 reads) and 71x (1,358,732,834 reads) respectively. To determine the location of the inserted gene cassette, we identified paired sequence reads for which one mate of the pair aligned to the rat genome (Rnor_6.0) and the other mate aligned to the inserted gene cassette. All such reads were aligned to the genome in the expected location in each line (24/24 D1-Cre, 25/25 A2a-Cre), indicating correct, single copy insertion (Figure 1C).

Partial sequence matches between the sgRNA and genomic locations away from the intended target may induce “off-target” cleavage events. Any off-target changes are likely to be progressively diluted over successive generations of back-crossing. We nonetheless performed an extensive screen and found no evidence for off-target events (see Methods).

### Consistent and specific Cre expression in *Drda1-expressing or Adora2a-expressing* cells

Our design ought to produce *Drd1a* (or *Adora2a*) mRNA in 1:1 stoichiometry with *iCre* mRNA, and thus yield highly selective *iCre* expression. To assess this we used triple fluorescent *in situ* hybridization, together with DAPI labeling of cell nuclei. Probe sets targeting *iCre, Adora2a* receptor and *Drd1a* receptor mRNA with distinct color labels were multiplexed and visualized simultaneously (Figure 2A). mRNA expression was quantified in three distinct striatal subregions - the dorsal striatum (DS), the nucleus accumbens core, and the nucleus accumbens medial shell. Automated software was used to define cell boundaries and count fluorescent puncta per cell, for each probe (Figure 2A).

**Figure 2.**
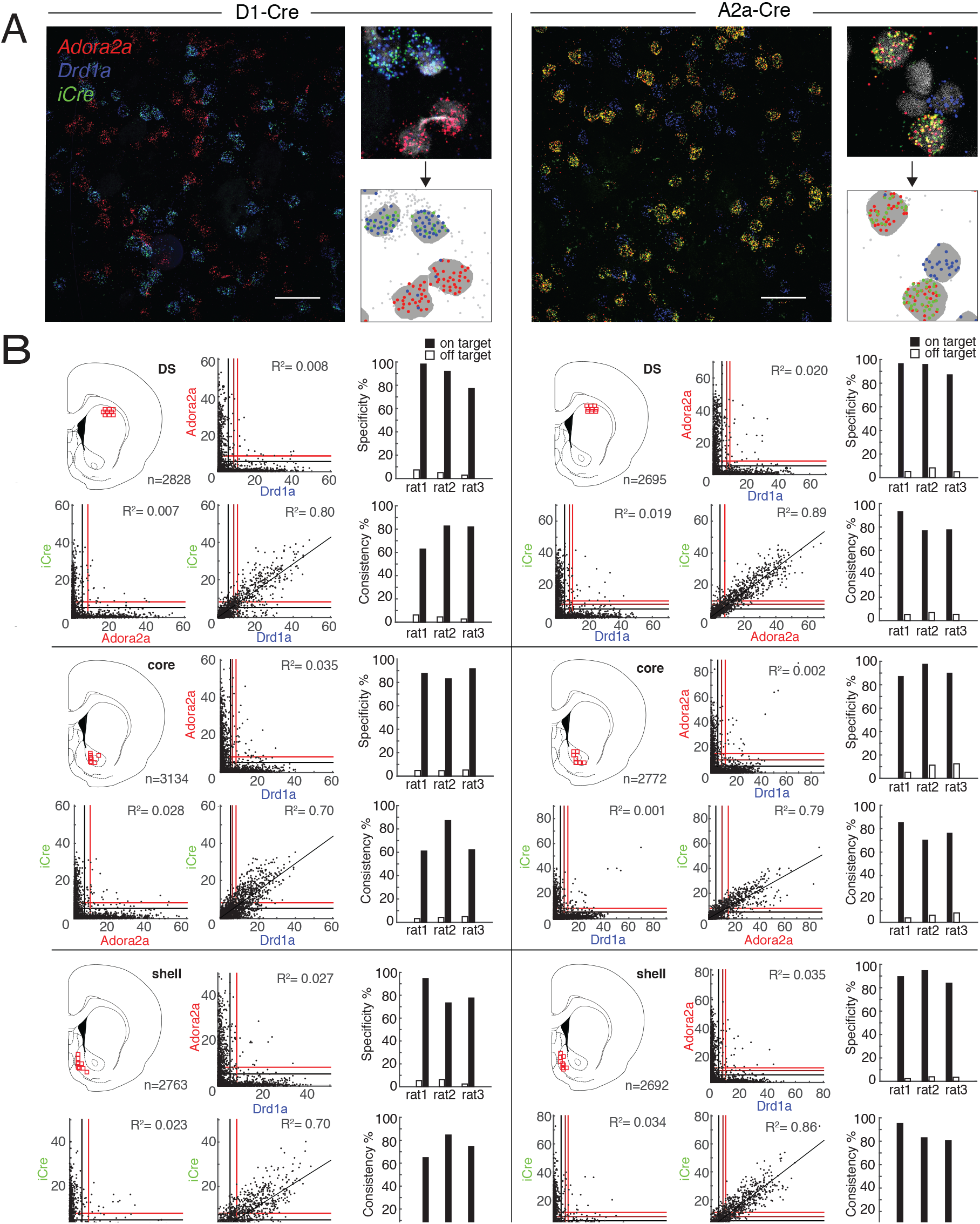
Confirmation and quantification of *iCre* production in D1+ and A2a+ MSNs. (A) Left of each column: example 40X images of FISH labelling used for quantification, taken from dorsal striatum (scale bars = 50 μm). Right of each column: closeup images (top) aligned with their corresponding automated software output (bottom). Gray regions indicate DAPI boundaries and colored dots indicate puncta within DAPI boundaries, using the same color scheme as the raw images. Gray dots indicate the locations of puncta detected outside of DAPI boundaries. (B) Scatterplots of raw puncta counts for each cell show selective *iCre* mRNA co-localization with the target receptor mRNA. Black, dark red and red lines indicate the 50^th^ (i.e. median), 95^th^ and 99.9^th^ confidence limits, respectively. Subpanels are grouped into rows by region and into columns by genotype. Atlas images depict the locations of confocal images used for mRNA quantification. Barplots show specificity and consistency of on-target and off-target expression, in each rat (n=3 rats per line).

Relationships between puncta counts for *Drd1a, Adora2a*, and *iCre* are shown in Figure 2B (left column, D1-Cre; right column, A2a-Cre). As expected, in D1-Cre rats expression of *Drd1a* and *iCre* mRNA was closely correlated in all striatal subregions examined (Fig. 2B; DS: R^2^=0.80; core: R^2^=0.70; shell: R^2^=0.70), and there was no correlation between *Adora2a* and *iCre* mRNA (DS: R^2^=0.008; core: R^2^=0.035; shell: R^2^=0.027). Conversely, in A2a-Cre rats expression of *Adora2a* and *iCre* mRNA was closely correlated (DS: R^2^=0.89; core: R^2^=0.79; shell: R^2^=0.86), and there was no correlation between *Drd1a* and *iCre* mRNA (DS: R^2^=0.019; core: R^2^=0.001; shell: R^2^=0.034). Consistent with earlier *in situ* hybridization studies (Berke et al., 1998; Le Moine and Bloch, 1995) we found near-complete segregation of dMSN and iMSN markers in all regions examined, with virtually no overlap between *Drd1a* and *Adora2a* expression (Figure 2A,B).

To further assess the specificity and consistency of *iCre* mRNA expression we defined thresholds for considering neurons as “positive” for a given probe. Given the wide distributions of puncta counts, the choice of threshold is non-trivial; it forces a tradeoff between type I and type II errors. Therefore, rather than picking an arbitrary threshold, for each probe we chose the 95% upper confidence limit, assuming a Poisson background distribution of puncta (see Methods). Using these thresholds (marked by red lines on the Fig.2B scatterplots) we estimated A2a-Cre specificity (% of *iCre*+ that are also *Adora2a*+) to be 93.5% (DS), 91.8% (core), and 89.2% (shell), and consistency (% of *Adora2a*+ that are also *iCre*+) to be 82.8% (DS), 77.4% (core), and 86.2% (shell). In the D1-Cre line, we estimated specificity (% of *iCre*+ that are also *Drd1a*+) to be 89.1% (DS), 87.4% (core), and 81.8% (shell), and consistency (% of *Drd1a*+ that are also *iCre*+) to be 77.5% (DS), 70.1% (core), and 74.6% (shell). If we use even higher thresholds for *Drd1a* and *Adora2a* (e.g. >30 puncta / cell) we can be essentially certain of cell identity, and assessed this way consistency was close to 100% for both lines (Figure 2B).

### Cre-dependent protein expression

We next examined whether *iCre* mRNA expression results in functional Cre protein confined to the appropriate basal ganglia pathway. To this end, we injected DS with a virus for Cre-dependent expression of a fluorescent protein (AAV-CAG-FLEX-tdTomato) and examined the expression pattern 4 weeks later. Consistent with pathway-specific expression of functional Cre protein, injection into the D1-Cre line resulted in clear expression in the striato-nigral pathway, while injection into the A2a-Cre line produced labeling in both DS and GPe, but no expression in the SNr (Figure 3A).

**Figure 3.**
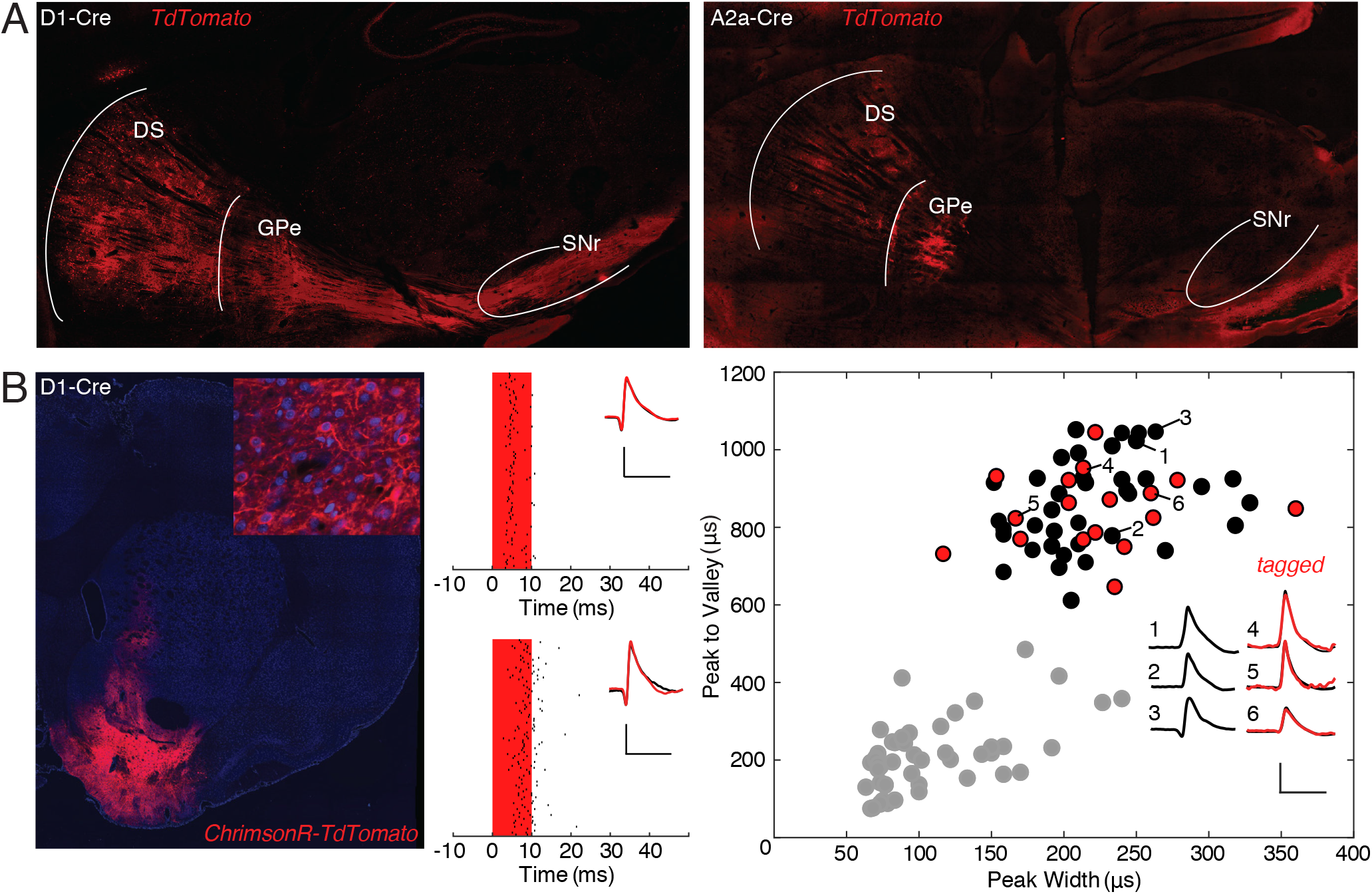
Cre-dependent expression confirms pathway segregation and functional expression. (A) Functional Cre expression is confined to appropriate BG pathways; (left) CAG-Flex-tdTomato injected into DS of the D1-Cre line expresses in terminals in SNr/GPi. (right) CAG-Flex-tdTomato injected into DS of the A2a-Cre line expresses in terminals in GPe but not SNr/GPi. (B) Optogenetic identification of Cre+ cells. (Left) hSyn-FLEX-ChrimsonR-TdTomato expression pattern into ventral striatum of a D1-Cre animal. Inset at top right shows closer-up view of transfected neurons. (Middle) Examples of a light-responsive neuron in a D1-Cre rat (top; identified dMSN) and an A2a-Cre rat (bottom; identified iMSN). Red bar indicates duration of light pulse, small black bars indicate spike times surrounding each stimulation (rows). Inset shows average session-wide spike waveform (black) with average light-evoked waveform overlaid in red. Scale bars, 0.1mV, 1ms. (Right) Waveform feature plot demonstrates that light-responsive dMSNs (red) are intermingled within the large cluster of presumed MSNs (black). Other, unclassified cells (mostly GABAergic interneurons) are shown in grey. Inset includes average spike waveforms from three examples each of light-responsive and non-responsive cells within the MSN cluster. Scale bars, 0.1mV, 1ms.

One important use of Cre lines is to enable positive identification of recorded neuron subtypes in awake behaving animals (e.g. Kravitz *et al*, 2010), via Cre-dependent opsin expression and monitoring neuronal responses to light pulses. We found that both the D1-Cre and A2a-Cre rat lines can be used for this purpose. In rats from each line we injected a virus (AAV-hSyn-FLEX-ChrimsonR-Tdtomato) into the accumbens core for Cre-dependent expression of the red-shifted opsin Chrimson (Klapoetke *et al*, 2014; Figure 3B, left) followed by a custom optrode (Mohebi et al., 2018). After allowing three weeks for opsin expression, we readily observed light-responsive single units (Figure 3B, middle). In a representative example session from a D1-Cre rat 17 neurons were identified as dMSNs, as they showed both a reliable response to red light stimulation and the waveform properties typical of MSNs (Figure 3B, right; Berke *et al*, 2004; Gage *et al*, 2010). As these cells were intermingled within the larger MSN cluster, it would not have been possible to identify them without this optogenetic tagging procedure.

### Normal acquisition and performance of instrumental and Pavlovian discrimination and cocaine-induced locomotor activity

Given that behavioral comparisons are likely to be made across these two independent transgenic lines, and between Cre+ rats and Cre-controls, we assessed acquisition and expression of instrumental responding for food and Pavlovian conditioned approach, and cocaine-induced locomotor activity in these lines.

In the instrumental discrimination task presses on an active lever were reinforced with food pellet delivery (fixed ratio of 1; FR1), whereas presses on an inactive lever were never reinforced. Rats were trained to an acquisition criterion of earning 50 pellets within less than 40 minutes. Figure 4A shows the average number of active and inactive lever responses, and Figure 4B depicts the average time to reach the acquisition criterion in each group. As expected active lever responding was greater than inactive lever responding and this did not differ between groups (Two-way repeated-measures ANOVA, main effect of lever: F_(1, 90)_ = 193.2, p<0.0001; n.s. main effect of group: F_(3, 90)_ = 1.379, p=0.2545; n.s. group x lever interaction: F_(3, 90)_ = 0.408, p=0.747). The time to reach acquisition criterion did not differ between groups (Two-way repeated-measures ANOVA, n.s. main effect of lineage: F_(1, 45)_ = 2.593, p=0.1143; n.s. main effect of genotype: F_(1, 45)_ = 1.578, p=0.2155; n.s. lineage x genotype interaction: F_(1,45)_ = 0.1086, p=0.7433).

**Figure 4.**
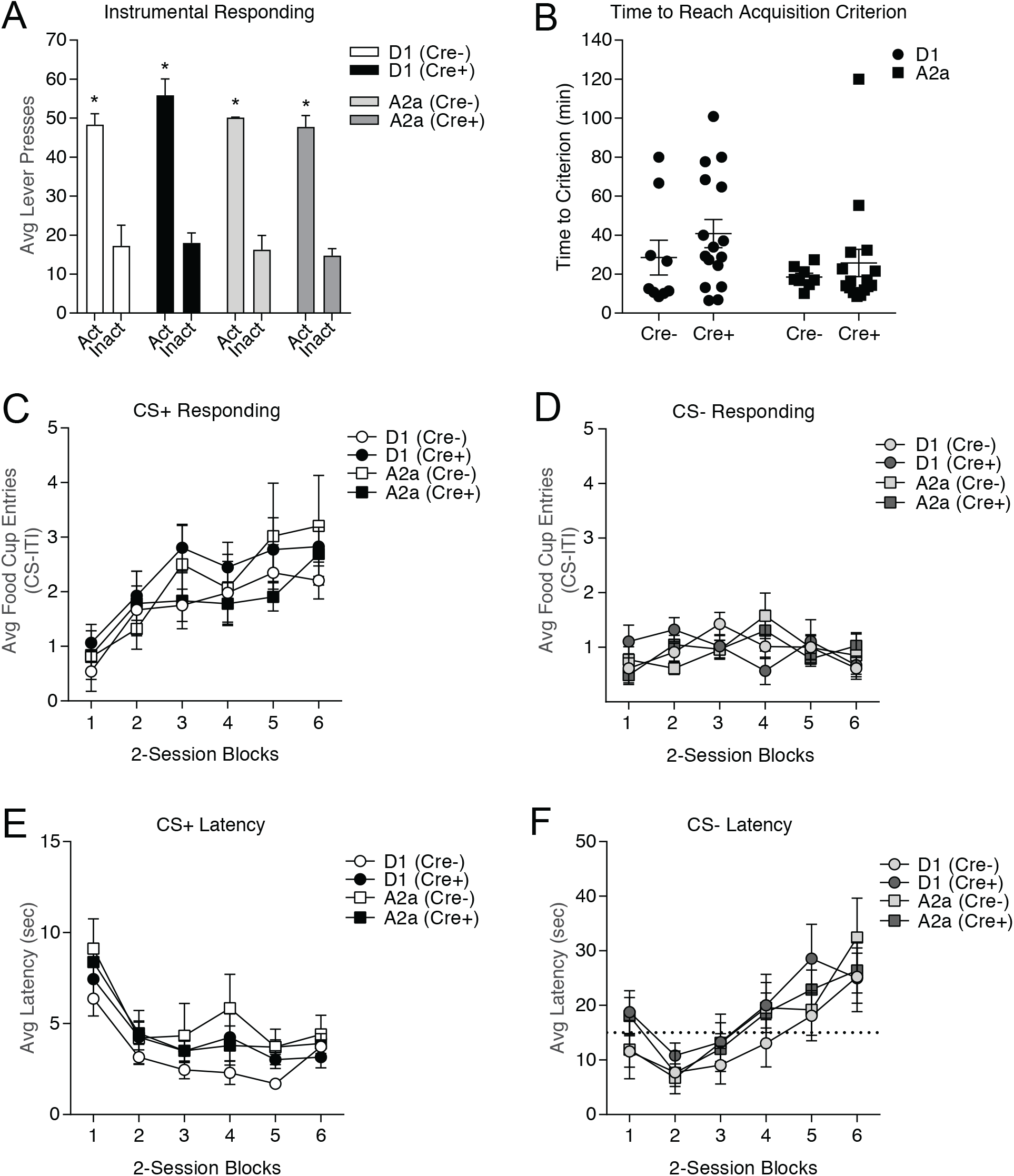
Instrumental and Pavlovian discrimination are similar between transgenic lines and Cre-littermate controls. (A) The average total number of responses on the active and inactive lever did not differ between groups and all groups preferentially responded on the active lever; *, p <0.05 active vs. inactive responses. (B) The total time to reach the acquisition criterion does not differ between groups. (C) The average rate of food cup entries during the first 10 seconds of CS+ presentations increases across 2-session training blocks and is similar between groups. (D) The average rate of food cup entries during the first 10 seconds of CS-presentations is low, does not change across training blocks and is similar between groups. (E) The average latency to approach the food cup following CS+ onset gets faster across training and is similar between groups. (F) The average latency to approach the food cup following CS-becomes slower across training and is similar between groups. Note the scale difference between panels E and F; the dotted line in panel F indicates 15 seconds on the y-axis to facilitate comparison. All data represented as mean ± SEM.

Following instrumental training, the acquisition and expression of Pavlovian conditioned approach were assessed in the same rats. During each session, one auditory cue was paired with food pellet delivery (conditioned stimulus: CS+), whereas a second auditory cue was never paired with food (CS-). Rats received 12 training sessions (60 min) in which each CS (tone or white noise, counterbalanced for CS+/CS-assignment) was randomly presented 4 times per session. Acquisition of Pavlovian conditioned food cup approach was similar across transgenic lines and between Cre- and Cre+ groups. Specifically, Figure 4C and D shows the average number of food cup entries during the first 10s of CS+ and CS-in 2-session blocks, respectively. Anticipatory food cup entries during CS+ presentations increased across training blocks and did not differ between groups (Figure 4C: Two-way repeated-measures ANOVA, main effect of training block: F_(5,225)_ = 13.45 p<0.0001; n.s. main effect of group: F_(3,45)_ = 0.505, p=0.6807; n.s. group x training block interaction: F_(15,225)_ =0.6086, p=0.866). In contrast, food cup entries during the first 10s of CS-presentations did not increase across sessions and was similar across groups (Figure 4D: Two-way repeated-measures ANOVA, n.s. main effect of training block: F_(5,225)_ = 1.602 p=0.1606; n.s. main effect of group: F_(3,45)_ = 0.01628, p=0.9971; n.s. group x training block interaction: F_(15,225)_ =1.476, p=0.1155). Thus, acquisition and maintenance of discriminatory conditioned approach were similar across transgenic lines, and between Cre- and Cre+ groups.

To provide an additional measure of learning we also examined the latency to enter the food cup following CS presentations. The average latency to enter the food cup following the onset of the CS+ decreased across training blocks and this decrease did not differ between groups, demonstrating that all groups were similarly motivated to respond to reward-predictive cues (Figure 4E: Two-way repeated-measures ANOVA, main effect of training block: F_(5,225)_ = 16.95, p<0.0001; n.s. main effect of group: F_(3,45)_ = 1.239, p=0.307; n.s. group x training block interaction: F_(15, 225)_ =0.3964, p=0.9791). In contrast, the average latency to enter the food cup following the onset of the CS-increased across training blocks, and did not differ between groups (Figure 4F: Two-way repeated-measures ANOVA, main effect of training block: F_(5,225)_ = 12.38, p< 0.0001; n.s. main effect of group: F_(3,45)_ = 0.6639, p=0.578; n.s. group x training block interaction: F_(15,225)_ =0.4812, p=0.9485). Together, the results from these behavioral studies show that introduction of Cre into either D1- or A2a neurons does not disrupt normal acquisition or expression of instrumental and Pavlovian discriminations.

Locomotor habituation and cocaine-induced locomotor activity were used to assess general striatal function in both lines (Oginsky et al., 2016). Cre+ rats and their Cre-littermates were placed in standard locomotor chambers equipped with photocell beams around the perimeter. After a 30 min habituation period, they were given two i.p. injections of saline (1 ml/kg). Both lines showed typical habituation to the locomotor chambers, and short-lived responses to saline injection that decreased with repeated injection. (Figure 5A, Two-way repeated-measures ANOVA, main effect of time: F_(21,378)_ = 10.42, p<0.0001). Locomotor activity was similar across D1-Cre and A2a-Cre lines and between Cre+ and Cre-rats (Figure 5A, Two-way repeated-measures ANOVA, n.s. effect of genotype: F_(2, 18)_ = 0.3965, p=0.6784). The next day, rats were again placed in locomotor chambers and given an injection of saline followed 40 min later by cocaine (15 mg/kg, i.p.). As expected, cocaine significantly increased locomotor activity, and the magnitude and time course of this response was similar between Cre+ and Cre-rats, as well as across transgenic lines (Figure 5B, Two-way repeated-measures ANOVA, main effect of time: F_(23,414)_ = 6.901, p<0.0001; n.s. effect of genotype: F_(2, 18)_ = 0.09284, p=0.9118; Figure 5C, Two-way repeated-measures ANOVA, main effect of injection: F(1,18) = 10.51, p=0.0045; n.s. effect of genotype: F(2, 18) = 0.1284, p=0.8803; n.s. injection by genotype interaction: F(2, 18) = 0.1122, p=0.8945). Thus, all genotypes showed a significant increase in locomotor activity following cocaine versus saline injection, and this effect did not differ between genotypes. These data suggest that there is no overt striatal dysfunction due to Cre expression, and that behavioral responses to elevations in dopamine are similar across D1-Cre and A2a-Cre lines.

**Figure 5.**
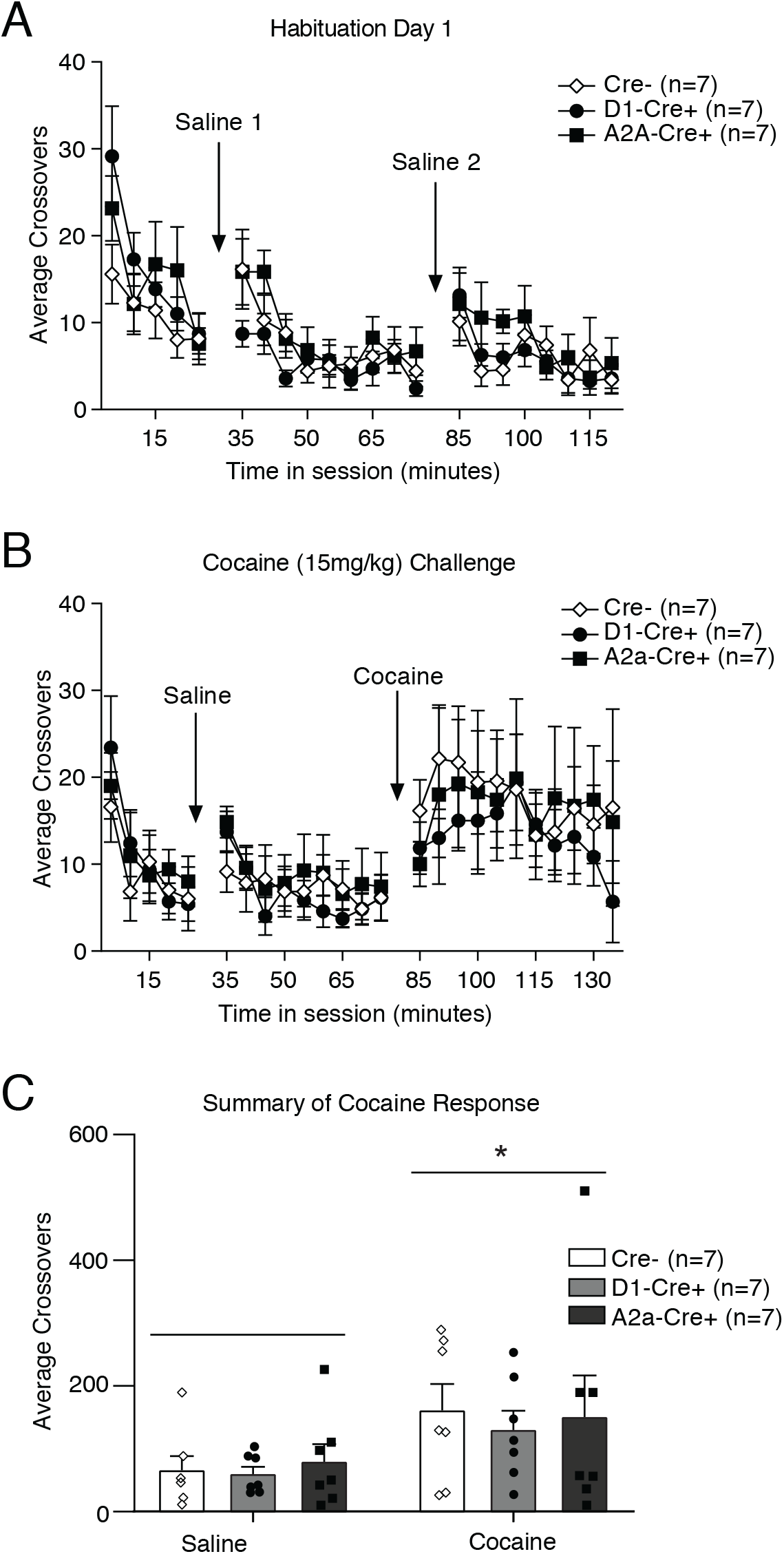
Basal and cocaine induced locomotor activity is similar between transgenic lines and Cre-littermate controls. (A) Locomotor activity decreases similarly in all groups across habituation and repeated saline injection. (B) Acute cocaine injection results in an increase in locomotor activity that is similar across groups. (C) Summary of locomotor activity in response to saline vs. cocaine. Cocaine significantly increases locomotor activity compared to saline, and the magnitude of this response is similar across groups; *, p<0.005 locomotor activity in response to cocaine vs saline. All data represented as mean ± SEM.

## DISCUSSION

We have demonstrated successfully-targeted, functional knock-in of Cre recombinase at the *Drd1a* and *Adora2a* loci, without off-target insertions as assessed by multiple methods including whole-genome sequencing. Comparable behavioral performance across lines and between Cre+ and Cre-littermates in several basic behavioral procedures provides further confidence that there are no deleterious effects of genetic manipulation or Cre expression. Within striatum we showed that Cre expression was consistent and selective to the correct populations of direct pathway D1+ and indirect pathway A2a+ cells respectively. Thus, these D1-Cre and A2a-Cre transgenic rats enable selective monitoring or manipulation of dMSNs and iMSNs with high specificity.

D1-Cre and A2a-Cre transgenic rats offer clear advantages over currently available transgenic models. First, the greater capacity of rats to learn complex behaviors make them stronger candidates for a wider range of tasks compared to mice. Second, the increased carrying capacity afforded by rats facilitates the chronic implantation of larger devices (i.e. high channel-count headstages, graded-refractive-index lenses). Thirdly, knock-ins can be used with higher confidence that the genetic modification was selective and specific to the target, compared to BAC lines.

A long-standing question in basal ganglia research has been the degree to which the striatal MSN population can be fully divided into distinct D1+ and D2+/A2a+ subpopulations. Based on BAC transgenic mice overlap has been reported to range from 4-5% in dorsal striatum and nucleus accumbens core, and up to 17% in shell (Bertran-Gonzalez et al., 2008; Wei et al., 2018). Our quantification of *Drd1* and *Adora2a* mRNA expression found overlap to be consistently very low in all striatal subregions examined, including shell, providing additional evidence for a fundamentally-segregated striatal architecture.

Since Cre expression was highly specific to the intended striatal pathways, these rats are powerful tools for pathway-specific neuron identification and manipulations. One caveat is that a subset of fast-spiking, parvalbumin-positive (PV+) interneurons also express D1 receptors (Bracci et al., 2002), and may thus also express Cre in D1-Cre rats. However, PV+ are only ~0.7% of striatal neurons (Luk and Sadikot, 2001) and at least in electrophysiological studies can be readily differentiated from MSNs (Berke, 2008; Kawaguchi, 1993; Koós and Tepper, 1999).

We chose to examine behavior during simple instrumental and Pavlovian tasks as well as cocaine-induced locomotor activity, as these behaviors rely heavily on striatal function. Although behavioral differences might emerge under other, more complex task conditions, the lack of any overt differences between the D1-Cre and A2a-Cre transgenic lines, or between Cre- and Cre+ littermates, strongly suggest that these rats are well-suited for behavioral and systems neuroscience studies. Beyond striatum, A2a receptors are found in the cortex, globus pallidus, hippocampus, thalamus, cerebellum (Rosin et al., 1998) and throughout the cardiovascular system. Similarly, D1 receptors are located in prefrontal cortex, hippocampus, thalamus and hypothalamus (Fremeau et al., 1991). In coordination with a rapidly expanding set of optical and genetic tools, these rats increase our ability to address fundamental questions about brain circuitry and mechanisms underlying neurological and psychiatric disorders.

### Funding and Disclosure

Support for this work was provided by National Institute on Drug Abuse awards R01DA045783 (JB), R21DA045277 (CF), T32DA007281 (RD) the National Institute on Neurological Disorders and Stroke award R01NS078435 (JB), the National Institute of Mental Health award R01MH101697 (JB), the National Institute of Diabetes and Digestive and Kidney Diseases awards R01DK106188 (CF), R01DK115526 (CF) and F31DK111194 (RD), the CHDI Foundation (JB), the University of Michigan, and the University of California, San Francisco. Support for the Transgenic Animal Model Core of the University of Michigan’s Biomedical Research Core Facilities was provided by The University of Michigan Cancer Center (NIH award P30CA46592, the University of Michigan Gut Peptide Research Center (NIH award P30DK34933) and the University of Michigan George M. O’Brien Renal Core Center (NIH award P30DK08194). Cocaine was provided by the NIDA drug supply program. The authors declare no conflicts of interest.

## Acknowledgements

TLS and JDB designed the *A2a-Cre and D1-Cre* constructs. EDH prepared the genome editing reagents. WEF performed rat zygote microinjections. JYY analyzed whole genome sequencing data and off-targets, and analyzed the FISH data with JRP. RCD performed and analyzed the behavioral experiments, which were supervised by CRF. TWF performed and analyzed *in vivo* electrophysiology. JDB conceived and supervised the project, and edited the manuscript. JRP performed the FISH and viral tracing experiments, and wrote the manuscript with input from all authors. We thank E. Wan and D. Vaka from the UCSF Institute for Human Genomics Core for performing the whole genome sequencing and assembly, H. Graham for assistance with FISH quantification, and R. Hashim, H. Bukhari, M. Zeidler, F. Ayres, Y. Alonso Caraballo, F. Sanchez Conde for assistance with breeding and genotyping.

The rat lines have been deposited with the Rat Resource and Research Center (rrrc.us) for community distribution (D1-Cre: RRRC#856, A2a-Cre: RRRC#857) and we thank E. Bryda for her help with this process.

